# MBOAT7-Driven Phosphatidylinositol Remodeling Promotes the Progression of Clear Cell Renal Carcinoma

**DOI:** 10.1101/648378

**Authors:** Chase K. A. Neumann, Renliang Zhang, Daniel J. Silver, Varadharajan Venkaleshwari, C. Alicia Traughber, Christopher Przybycin, Defne Bayik, Jonathan D. Smith, Justin D. Lathia, Brian I. Rini, J. Mark Brown

## Abstract

The most common kidney cancer, clear cell Renal Cell Carcinoma (ccRCC) is closely associated with obesity. In fact, the “clear cell” variant of RCC is given this name due to large lipid droplets within the tumor cells. Although it is well appreciated that renal lipid metabolism is altered in ccRCC, the mechanisms driving this are not well understood. Leveraging a shotgun lipidomics approach we have identified a lipid signature for ccRCC that includes an increase in arachidonic acid-enriched phosphatidylinositols (AA-PI). In parallel, we found that ccRCC tumors have increased expression of the acyltransferase enzyme membrane bound O-acyltransferase domain containing 7 (MBOAT7) that contributes to AA-PI synthesis. In ccRCC patients, *MBOAT7* expression increases with tumor grade, and increased *MBOAT7* expression correlates with poor survival. Genetic deletion of MBOAT7 in ccRCC cells decreases proliferation and induces cell cycle arrest, and MBOAT7^−/−^ cells fail to form tumors *in vivo*. RNAseq of MBOAT7^−/−^ cells identified alterations in cell migration and extracellular matrix organization, which were functionally validated in migration assays. Our work highlights the accumulation of AA-PI in ccRCC and demonstrate a novel way to decrease the AA-PI pool in ccRCC by limiting MBOAT7. Our data reveal that metastatic ccRCC is associated with altered AA-PI metabolism, and identify MBOAT7 as a novel target in advanced ccRCC.

## Introduction

The most lethal and third most common urological cancer in the United States is clear cell renal cell carcinoma (ccRCC) (1). Incident risk of ccRCC is closely associated with obesity (2), and ccRCC tumors are strikingly lipid laden compared to other malignancies (3). In fact, the “clear cell” variant of RCC is given this name due to large intracellular lipid droplets that accumulate within the tumor cells, and in some cases ccRCC cells have a unilocular lipid droplet similar to what is seen in adipocytes (4). Recently, several studies have demonstrated that altered prostaglandin levels may be involved in ccRCC growth and progression (5–7), and intracellular lipid droplets may play a role in managing oxidative stress in the ccRCC tumor microenvironment (4). Although it is well appreciated that renal lipid metabolism is altered in ccRCC, the mechanisms driving this are not well understood.

Phosphatidylinositols (PI) and closely related phosphoinositides (PIPs) are essential for life in eukaryotes (5). These signaling phospholipids are commonly studied for their importance in cytokinesis, migration and cell polarity (6–8). Although many studies have identified key signaling roles for PIPs in cancer progression (8–10) very little is known regarding whether compositional changes in the membrane PI pool, where PIP signaling lipids originate, is involved in ccRCC pathogenesis. Using acyl-CoAs as donors, phospholipids are formed by the *de novo* “Kennedy” pathway (12) and diversified by a remodeling pathway known as the Lands’ cycle (13). Although the major enzymes involved in the Kennedy pathway have been well characterized (12), much less is known about enzymes involved in the Lands’ cycle. A unique contributor to the membrane PI pool is membrane bound O-acyl transferase domain containing 7 (MBOAT7), an acyltransferase enzyme that selectively esterifies lysophosphatidylinositol (LPI) lipids to an arachidonyl-CoA to form the major PI species (38:4) in the inner leaflet of cell membranes (14, 15). MBOAT7 is a unique contributor to the Lands’ cycle, which is a series of phospholipase-driven deacylation and lysophospholipid acyltransferase-driven reacylation reactions that synergize to alter phospholipid fatty acid compositions, creating membrane asymmetry and diversity. It is important to note that MBOAT7, unlike other lysophospholipid acyltransferses, only diversifies the fatty acid composition of membrane PI species and not phospholipids with other head groups (15, 16) Here we demonstrate that differential expression of MBOAT7 in advanced ccRCC is associated with survival, and MBOAT7 loss of function diminishes both proliferative and migratory properties of ccRCC cells. These data support the notion that limiting MBOAT7-driven PI diversification may hold therapeutic promise in ccRCC and potentially other cancers.

## Results

### MBOAT7 and its enzymatic product PI 36:4 are increased in ccRCC tumors

To study the possible lipid related drivers of ccRCC, we took primary tumor biopsies with matched normal adjacent tissue samples from patients at the Cleveland Clinic, and performed untargeted lipidomics **(Figure 1A)**. One of the most significantly increased lipids in the tumors compared to the non-tumor was PI 36:4 **(Figure 1B)**. In parallel, expression of the LPI acyltransferase enzyme *MBOAT7* was elevated **(Figure 1C)**. Previous literature has demonstrated that MBOAT7 is the primary LPI acyltransferase enzyme involved in the enrichment of arachidonic acid into PI (AA-PI) (17) within the Lands’ cycle of phospholipid remodeling **(Figure 1D)**.

**Figure 1.**
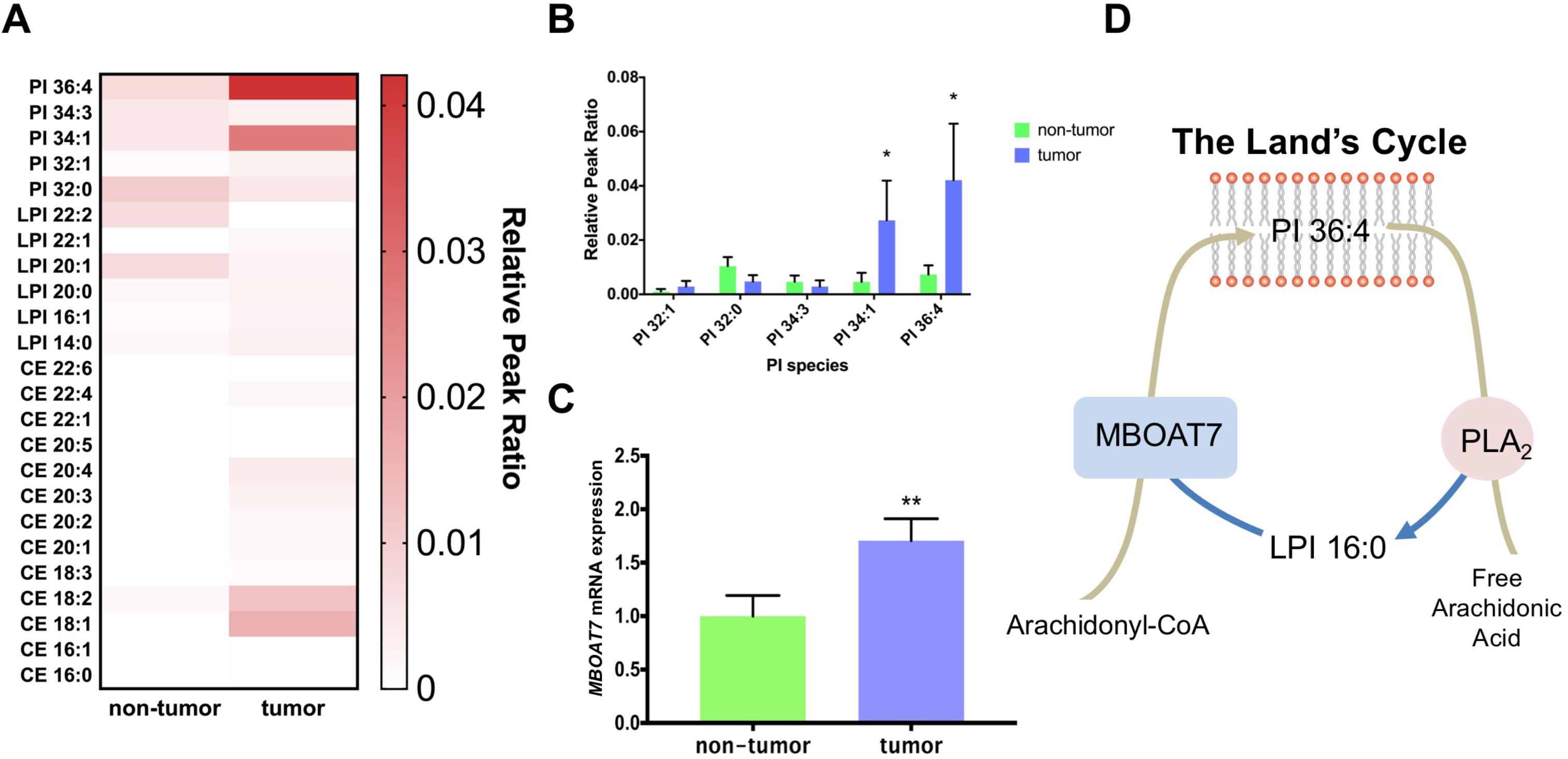
Arachidonic acid containing-PI (AA-PI) and MBOAT7 are elevated in ccRCC tumors. **(A)** Heatmap of most significantly altered lipids in untargeted lipidomic profiling of metastatic ccRCC tumor samples and matched non-tumor samples; n=10 per group. **(B)** Phosphatidylinositol (PI) species in the tumor and non-tumor samples; n=10 per group. **(C)** mRNA levels of the MBOAT7, a key acyltransferase determining AA-PI levels. **(D)** Schematic representation of the Lands’ cycle of PI remodeling. All data are presented as mean ± S.E.M., unless otherwise noted. Student t-test: * < 0.05, ** < 0.0021, *** < 0.0002

### MBOAT7 increases with ccRCC severity and correlates with poor survival

To explore the association of *MBOAT7* expression within histological grade of ccRCC, we analyzed nephrectomy samples from initial surgical resection with age-, BMI-, and sex-matched patients. Interestingly, *MBOAT7* expression increases with histological grade of ccRCC, particularly in grade IV cases **(Figure 2A)**. Furthermore, a commonly used ccRCC cell line Caki-1 has an increased expression of *MBOAT7* similar to that of the grade 4 tumor samples **(Supplementary Fig. 2)**. We next confirmed the increased *MBOAT7* expression with histological grade in a validation cohort of patient samples. The Cancer Genome Atlas (TCGA) ccRCC cohort (KIRC) demonstrated a similar finding that *MBOAT7* expression increases with histological severity **(Figure 2B)**. To assess the relationship between *MBOAT7* mRNA expression and patient survival, we leveraged access to TCGA datasets and stratified patients into low versus high *MBOAT7* mRNA levels. High *MBOAT7* expression correlates with a poorer overall survival within histology matched cohorts or Stage III/IV patients **(Figure 2C)**. These data provide evidence that *MBOAT7* and enzymatic substrates or products may play a role in progression of ccRCC.

**Figure 2.**
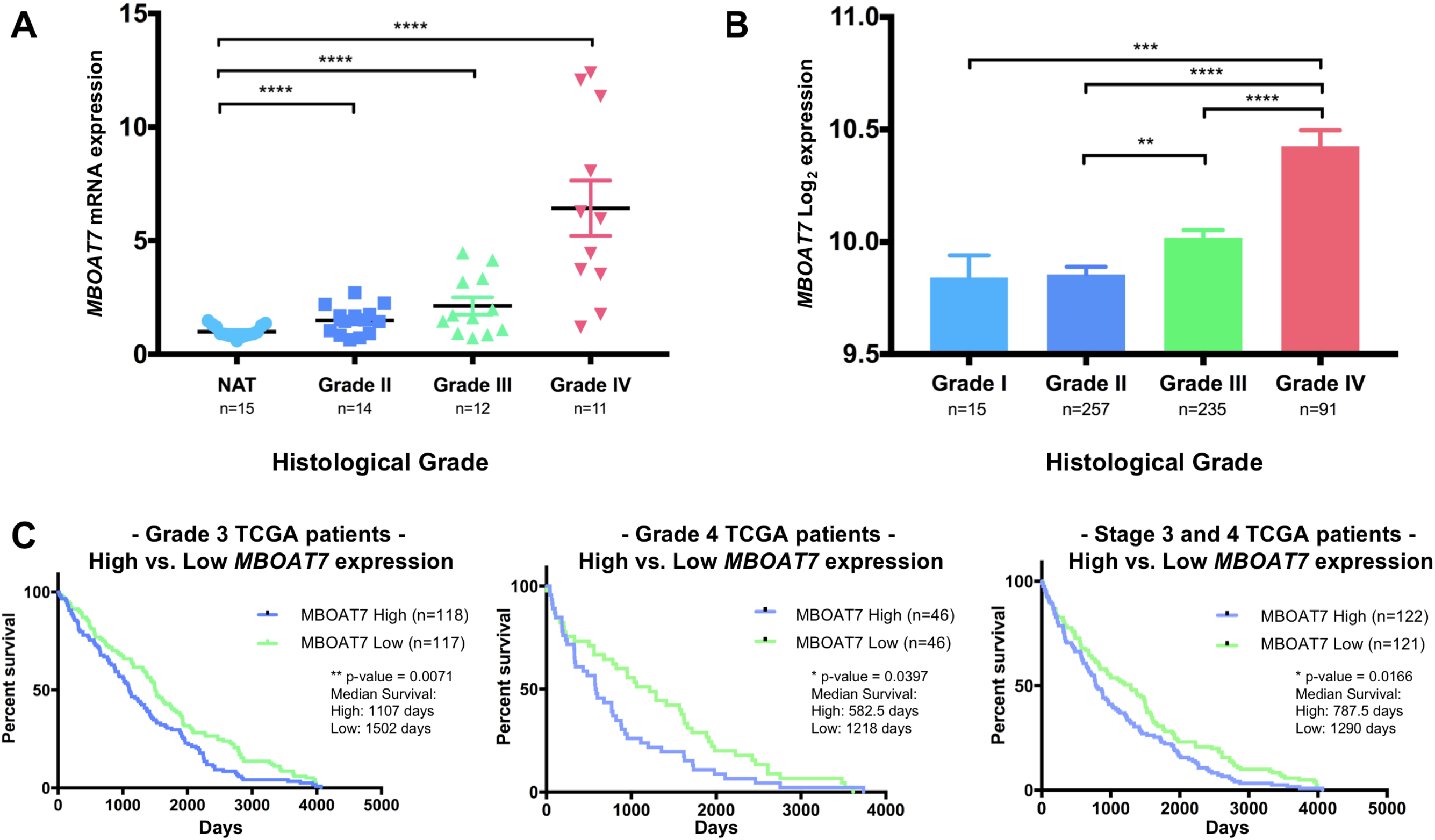
MBOAT7 expression increases with ccRCC severity and correlates with poorer survival. **(A)** MBOAT7 mRNA expression in ccRCC patients with increasing severity of histological grade in patient tumor biopsies. NAT = Normal Adjacent Tissue. **(B)** A validation dataset from the cancer genome atlas (TCGA) stratified by pathological staging (I-IV) and pan-cancer normalized. **(C)** High MBOAT7 expression significantly correlates with a decrease in overall survival in TCGA patients stratified by Grade 3 and Grade 4 Histological neoplasms. Similarly, High MBOAT7 expression correlates to a decrease in overall survival with Stage III and IV patients. One-way Annova and Logrank Test (Mantel-Cox test) ns > 0.1234, * < 0.05, ** < 0.002, *** < 0.0002

### Generation and validation of MBOAT7 loss of function in ccRCC cell line

To assess the role of *MBOAT7* in ccRCC, we utilized CRISPR/Cas9 to knockout MBOAT7 in the commonly used ccRCC cell line Caki-1. Following clonal isolation, we confirmed knockout using sequencing across exon 6 **(Supplementary Fig. 2)** as well as quantification of mRNA and protein abundance **(Figure 3A)**. We then validated this loss of enzymatic function with targeted mass spectrometry of MBOAT7 LPI substrates and PI product species in targeted cells. Recall that MBOAT7 acts on LPI 16:0 to generate a series of AA-PI species including PI 36:4. Thus, as a result of MBOAT7 deletion, the levels of LPI substrates are significantly increased in both MBOAT7^−/−^ clones **(Figure 3B)** and the set of AA-PI (36:4, 38:4, 38:5) products were significantly attenuated **(Figure 3C)**. The functional changes resulting from MBOAT7 knockout were finally validated by providing exogenous ^3^H-aracidonic acid to MBOAT7 null cells. Both MBOAT7^−/−^ clones exhibited a ~90% reduction in AA incorporation into PI species **(Supplementary Fig. 3)**.

**Figure 3:**
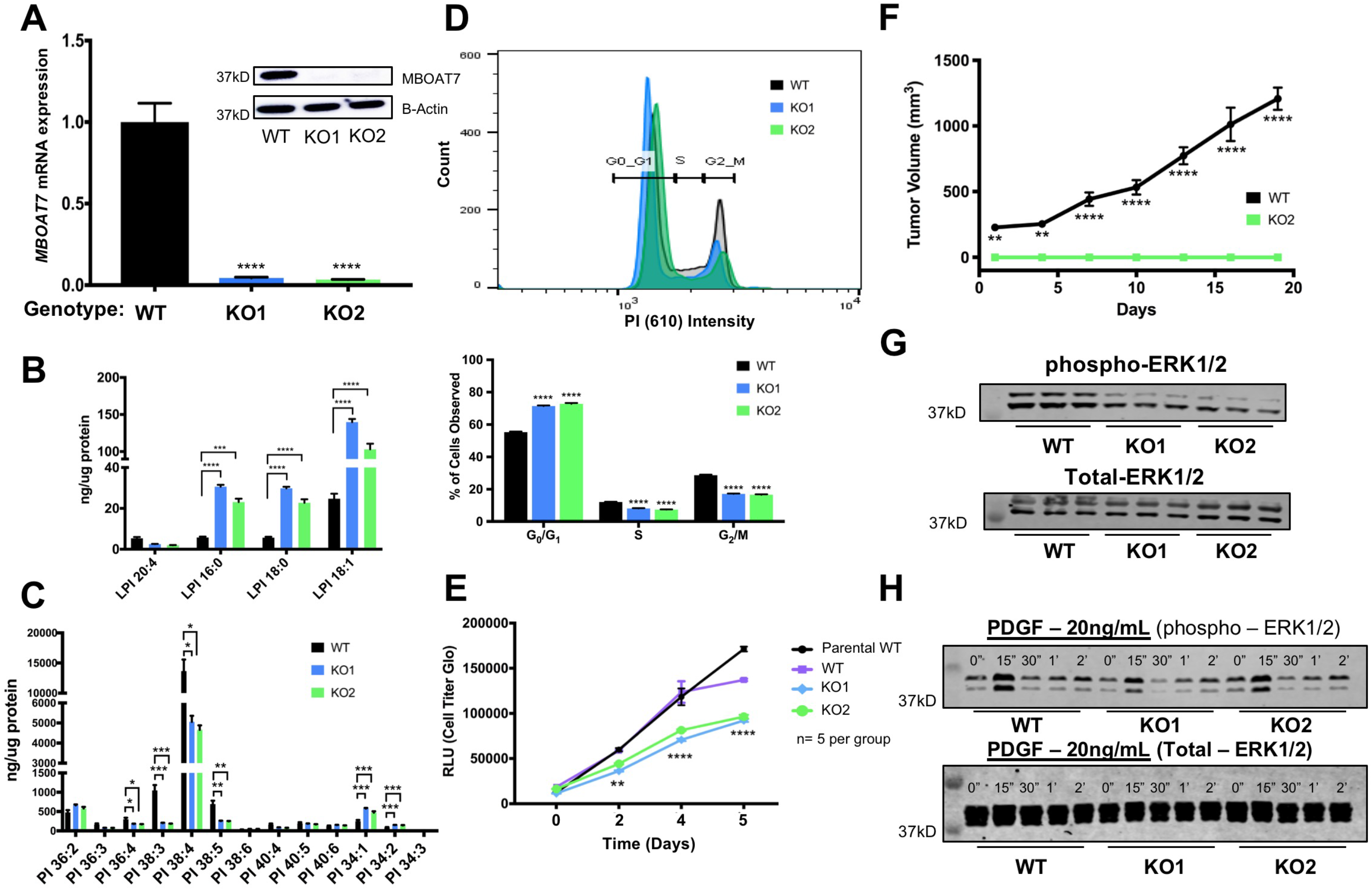
Genetic deletion of MBOAT7 in ccRCC results in reduced arachidonic acid-PI, cell cycle, proliferation, and growth factor signaling. **(A)** Generation of Caki-1 MBOAT7 knockout (KO) cell lines: KO1 and KO2. **(B)** Targeted mass spectrometry of Caki-1 WT, KO1, and KO2 cell lines show increased LPI substrate in MBOAT7 KO1 and KO2. **(C)** Targeted PI mass spectrometry of Caki-1 WT, KO1, and KO2 showed significantly decreased arachidonic acid containing-PI (AA-PI). **(D)** Cell cycle analysis in WT, KO1, and KO2 cell lines (n = 4 per group). **(E)** Proliferation assay (Cell Titer Glo) of Caki-1 MBOAT7 KO cell lines, WT, and parental Caki-1 plated at 2000 cells/well at Day 0. Confirmed in three independent experiments. **(F)** *In Vivo* xenograft model with subcutaneous injection of MBOAT7 WT vs. KO2 tumor growth show decreased engraftment of MBOAT7 KO *in vivo.* **(G)** Western Blot analysis of Asynchronous signaling via phospho-ERK1/2 and Total-ERK1/2. (**H)** Serum-starved synchronized PDGF growth factor time course. All results shown are representative from one experiment that was replicated in at least 2-3 independent experiments. Multiple and Student t-test with p-value significance: ns > 0.1234, * < 0.05, ** < 0.002, *** < 0.0002, **** < 0.0001

### Loss of function MBOAT7 is associated with decreased cell proliferation, cell cycle, and in vivo tumor formation

To assess phenotypic changes in MBOAT7^+/+^ and MBOAT7^−/−^, we pursued cell proliferation, cell cycle, and subcutaneous xenograft growth experiments. When cell cycle was analyzed MBOAT7^−/−^ cells show an increased number of cells in G_0_/G_1_ while decreased S and G_2_/M phases **(Figure 3D)**. In parallel, MBOAT7^−/−^ cells showed significant reduction in proliferation up to ~35% over 5 days **(Figure 3E)**. To test whether this had any effect on tumor growth *in vivo,* we performed in vivo xenograft experiments. Approximately 8 weeks post injection, the MBOAT7^+/+^ cells had reached endpoint in the tumor growth experiment, but the MBOAT7^−/−^ failed to develop a palpable tumor **(Figure 3F)**. A common growth-signaling pathway related to ccRCC cellular proliferation is the mitogen-activated protein kinase (MAPK) cascade (18–20). In an asynchronous culture, the MBOAT7^−/−^ cells have a reduced phospho-ERK1/2 levels **(Figure 3G)**. Upon synchronizing cells, the MBOAT7^−/−^ cells demonstrate a blunted platelet-derived growth factor stimulated response in phospho-ERK1/2 activation **(Figure 3H)**.

### MBOAT7^−/−^ ccRCC cells shows differentially regulated motility, proliferation, and matrix organization

We performed an unbiased RNAseq of MBOAT7^+/+^ and MBOAT7^−/−^ Caki-1 cells. The principal component analyses (PCA) analysis of both knockout populations demonstrates majority of the variation can be defined by the loss of MBOAT7 function **(Supplementary Fig.4)**. This unbiased analysis demonstrated a group of genes conserved and differentially regulated among both knockout populations compared to wild-type **(Figure 4A)**. Upon pathway analysis of both knockouts, the pathways revealed a conserved significant enrichment in NABA ECM Glycoproteins, negative regulation of cell differentiation, negative regulation of cell migration, and negative regulation of cell proliferation **(Figure 4B)**. To validate this RNAseq dataset, we performed RT-PCR on four up-regulated (*SOX6*, *CDH6, FBN1, and ADAMTS5)* and four down-regulated (*ACSL5, CPT1C, FN1, and KLHL14)* transcripts **(Figure 4C)**. Functionally, to follow-up the RNAseq pathway results, we pursued an *in vitro* migration assay to assess the potential differences in motility.

**Figure 4:**
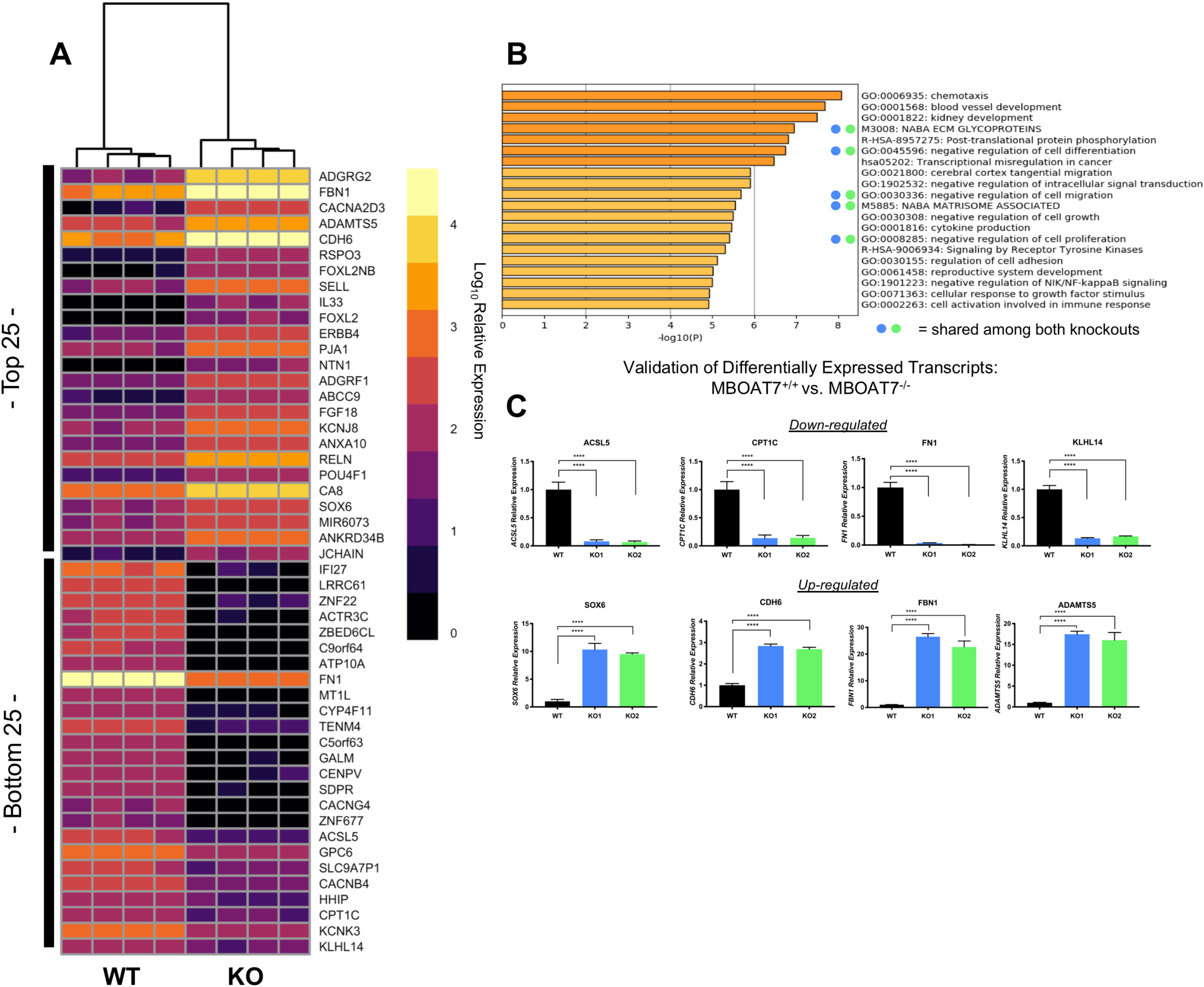
RNA sequencing in MBOAT7 deficient ccRCC cells reveals alterations in cell cycle, matrix organization, and cell migration. **(A)** Heatmap of Top 25 and Bottom 25 differentially expressed transcripts from unbiased RNAseq. **(B)** Pathway analysis reveals regulation of migration, cell proliferation and extracellular organization from top 250 significantly upregulated transcripts. **(C)** qPCR validation of selected genes altered in MBOAT7 null Caki-1 cells. Student t-test and p-value: * < 0.05, ** < 0.002, *** < 0.0002, **** < 0.0001

### MBOAT7^−/−^ ccRCC cells show decreased migration and less mesenchymal phenotype

Our RNAseq pathway analysis led to multiple implications suggesting decreased migration and matrix organization. Using an *in vitro* scratch assay, we found that MBOAT7^−/−^ clones demonstrates a ~50% reduction in motility **(Figure 5B)**. Videos of the MBOAT7 loss of function also demonstrates a change in phenotype while migrating **(Supplemental Movie 1-3)**. To further explore potential differences in motility using qPCR, we looked at the expression of specific mesenchymal and epithelial genes: *SNAIL, ZEB1*, *ACTA2*, and *CDH1.* The MBOAT7^−/−^ have decreased expression of mesenchymal genes: *SNAIL*, *ZEB1*, *ACTA2*, yet have increased expression of the epithelial marker E-cadherin (*CDH1)* **(Figure 5C)**. At the protein level, a similar loss of mesenchymal markers and increased epithelial markers has occurred in our MBOAT7^−/−^ cells **(Figure 5D)**. These data suggest that MBOAT7-driven phospholipid remodeling is a critical determinate of epithelial to mesenchymal transition (EMT).

**Figure 5:**
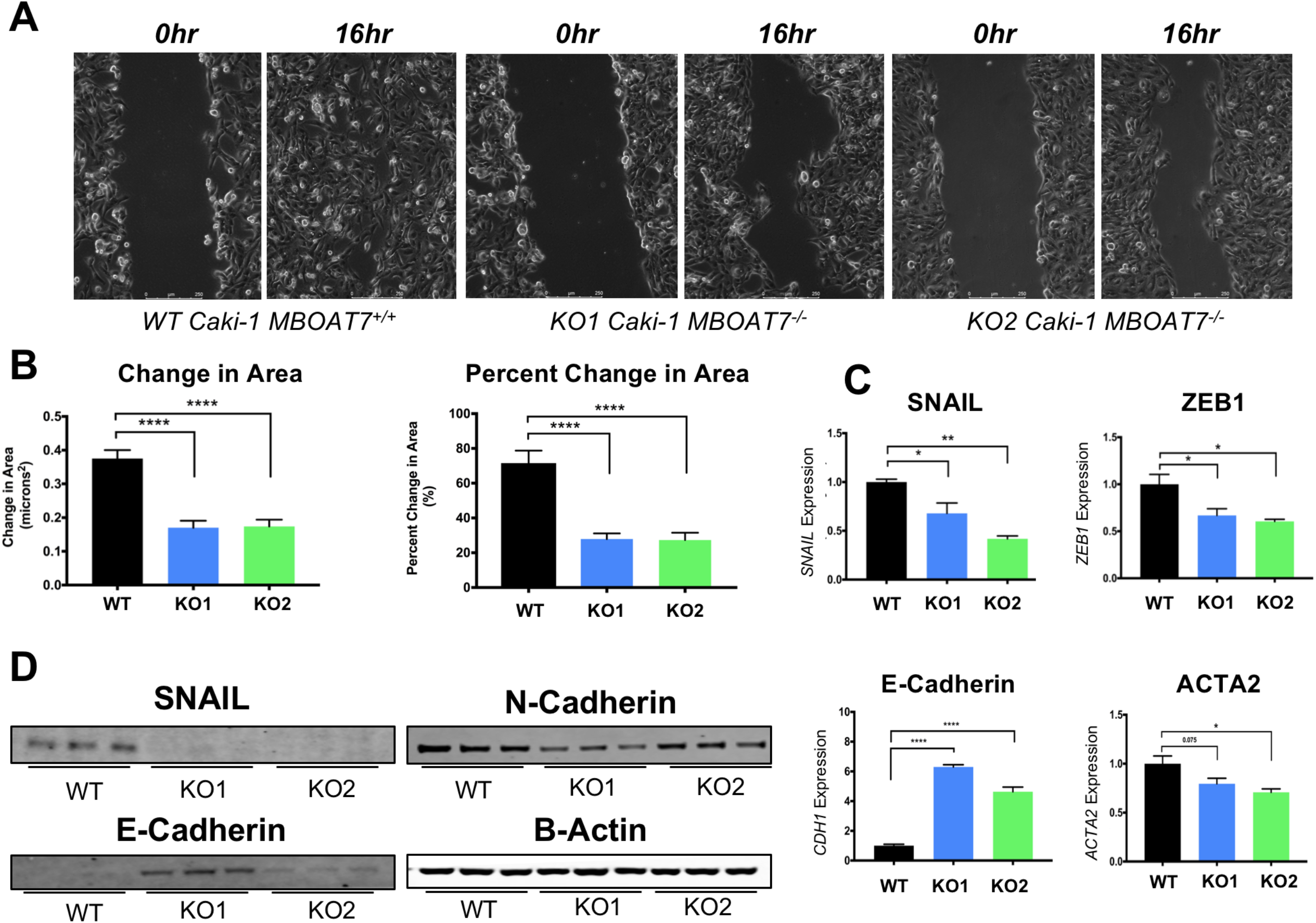
MBOAT7 regulates in vitro migration and epithelial/mesenchymal markers. **(A)** Representative micrographs of MBOAT7^+/+^ and MBOAT7^−/−^ cells showing in vitro scratch assay (time 0 and time 16hr); See online supplement for time lapse images. **(B)** Quantification of change in area and percent change in area in the *in vitro* scratches. **(C)** qPCR of MBOAT7^+/+^ and MBOAT7^−/−^ demonstrates altered expression of epithelial and mesenchymal marker genes. **(D)** Western blot analysis shows a decrease in mesenchymal markers and increase in epithelial markers with the loss of MBOAT7. All results shown are representative from one experiment that was replicated in at least 2-3 independent experiments (n=2-5 per group). Student t-test and p-value: * < 0.05, ** < 0.002, *** < 0.0002, **** < 0.0001

## Discussion

Similar to other solid tumors, ccRCC is primarily driven by loss of key tumor suppressor such as von Hippel-Lindau (VHL) and activation of oncogenes (23–25). Closely associated with these genetic alterations, lipid metabolic processes are dramatically altered in the ccRCC tumor microenvironment (3, 4, 26–33). In fact, the “clear cell” subtype gets its name from cholesterol ester-rich lipid droplets that accumulate in tumor cells (32, 33). Although it has long been appreciated that ccRCC can be characterized as a “lipid metabolic disease” (34–36), there is still incomplete understanding of which lipid metabolic targets are therapeutically tractable. This study identifies MBOAT7-driven PI remodeling as a new potential drug target. The key findings from the current study are: (1) In addition to accumulation of cholesteryl ester (32, 33), metastatic ccRCC is characterized by accumulation of AA-PI lipids and elevated expression of the AA-PI synthesizing enzyme MBOAT7, (2) MBOAT7 expression is elevated in high grade ccRCC, and high MBOAT7 expression is associated with poor survival, (3) Genetic deletion of MBOAT7 in ccRCC cells results in reduced AA-PI levels and reciprocal increases in substrate LPIs, (4) MBOAT7 null ccRCC cells exhibit reduced proliferation associated with cell cycle arrest, and fail to form tumors *in vivo*, (5) MBOAT7 null ccRCC cells have reduced growth factor-driven MAPK activation, (6) Unbiased RNA sequencing also reveals a role for MBOAT7 as a key regulator of ccRCC migration and epithelial-to-mesenchymal transition (EMT). Collectively, our findings identify a key new lipid metabolic dependency in ccRCC, and show that inhibition of MBOAT7-driven remodeling of membrane PI lipids may hold therapeutic potential in patients suffering from this devastating disease.

The closely related signaling network of phosphoinositide 3-kinase (PI3K), protein kinase B (AKT), and mammalian target of rapamycin (mTOR) controls almost all aspects of cancer initiation and progression. Given the central role that the PI3K-AKT-mTOR pathway plays in cancer cell biology, many developmental and approved cancer drugs target key nodes within the pathway (37–39). Although many drugs currently target the PI3K-AKT-mTOR pathway are designed towards kinase inhibition, it is important to remember that signaling through the PI3K-AKT-mTOR pathway is sustained by metabolism of key lipids such as PI (9, 40–43). The PI pool in the inner leaflet of the plasma membrane is quantitatively small compared to other phospholipids, but serves as a key reservoir for growth factor and hormone stimulated generation of phosphatidylinositol phosphate (PIP) lipids (44–46). Key PI-derived lipids such as phosphatidylinositol 4-phosphate (PI4P), phosphatidylinositol 4,5-bisphosphate (PI-4,5-P_2_), and inositol 1,4,5-triphosphate (IP_3_) play critical roles in normal hormone and growth factor signaling and aberrant signaling processes that drive malignancy (44, 45, 47, 48), and as a result many anti-cancer drugs aim to alter levels of these critical signaling lipids. However, very little attention has been paid to factors that regulate the abundance or composition of the membrane PI pool where PIPs originate. Here, we have identified a lipid signature of ccRCC that is characterized by specific alterations in arachidonic acid-containing membrane PI species (AA-PI), which is likely to have downstream regulatory roles in PIP-dependent signaling. In support of this concept, it was recently demonstrated that MBOAT7 knockout mice have a selective reduction in arachidonic acid-containing PIP species (15). The decrease of the PI pool may have a broader impact to cell function then PIP_3_ formation, as this may also impact the vesicular trafficking. The decreased response to growth factor stimulation, may be due to the decreased PtdIns(4)P in the endosomal compartment important for growth factor receptor sorting (21, 22). Currently, it is incompletely understood how the acyl chain composition of PIP lipids can impact downstream kinase activation, but our studies suggest that this is an area of research worth exploring across many diverse cancers.

Given the striking lipid accumulation seen in ccRCC, it remains entirely possible that abnormal lipid metabolism in the tumor microenvironment is mechanistically involved in tumor initiation and/or progression. However, the challenge that lies ahead is to identify the key lipid metabolic targets that are therapeutically tractable. Some of the earliest lipidomic studies identified striking accumulation of cholesterol esters in cytosolic lipid droplets of ccRCC patients (3, 33). However, it is unlikely this inert storage form of lipid plays a causal role in ccRCC pathogenesis, and instead likely is a result of altered cholesterol handling in transformed cells. A more recent study suggested that esterification of fatty acids into triglyceride may provide a “buffer” for the toxic effects of saturated ceramides and acylcarnitines (30). However, in contrast to the large accumulation of cholesterol ester, human ccRCC tumors are not particularly rich in stored triglyceride demonstrating that this “buffering” capacity is likely limited. There is also evidence that hypoxia-inducible factor (HIF) driven transcriptional repression of carnitine palmitoyltransferase (CPT1A) plays a role in ccRCC tumorigenesis, but there are significant hurdles in trying to re-express a silenced gene in cancer cells. Furthermore, HIF2α hyperactivity in ccRCC cells can promote the expression of the lipid droplet-associated protein perilipin 2 (PLIN2), which facilitates lipid storage and indirectly blunts endoplasmic reticulum (ER) stress pathways (4). It is interesting to note that PLIN2 has been identified as a potential urinary biomarker of advance ccRCC (48, 49), providing additional rationale to develop therapeutic strategies to intervene on this HIF2α-PLIN2 pathway. Collectively, all of these recently described lipid metabolic pathways altered in ccRCC provide potential targets for drug discovery. Here we expand this list to include MBOAT7 as a potential lipid metabolic target that is therapeutically tractable using a number of platforms. Currently, there is a semi-selective small molecule MBOAT7 inhibitor (thimerosal), (50) which has shown some promise as a potential as a cancer therapeutic (51–53). However, thimerosal likely has many enzymatic targets, and additional structure-activity-relationship (SAR) drug discovery efforts are needed to find more selective inhibitors. Collectively, our work suggests that selective MBOAT7 inhibitors may hold promise to blunt the progression of metastatic ccRCC, and these findings may be broadly related to other cancer types where the PI3K-AKT-mTOR axis is hyperactive.

## Methods

### Lipidomic Profiling of ccRCC

To understand alterations across a wide range of structurally-distinct lipids we developed a shotgun lipidomics method for measurement of multiple lipid species as previously described (54). All the internal standards were purchased from Avanti Polar Lipids, Inc. (700 Industrial Park Drive, Alabaster, Alabama 35007, USA). Ten internal standards (12:0 diacylglycerol, 14:1 monoacylglycerol, 17:0 lysophosphatidylcholine, 17:0 phosphatidylcholine, 17:0 phosphatidic acid; 17:0 phosphatidylethanolamine, 17:0 phosphatidylglycerol, 17:0 sphingomyelin, 17:1 lysosphingomyelin, and 17:0 ceramide) were mixed together with the final concentration of 100 μM each. Total hepatic lipids were extracted using the method of Bligh and Dyer (Bligh and Dyer 1959) with minor modifications. In brief, 50 μL of 100 μM internal standards were added to tissue homogenates and lipids were extracted by adding by adding MeOH/CHCl3 (v/v, 2/1) in the presence of dibutylhydroxytoluene (BHT) to limit oxidation. The CHCl_3_ layer was collected and dried under N_2_ flow. The dried lipid extract was dissolved in 1 ml the MeOH/CHCl_3_ (v/v, 2/1) containing 5mM ammonium acetate for injection. The solution containing the lipid extract was pumped into the TripleTOF 5600 mass spectrometer (AB Sciex LLC, 500 Old Connecticut Path, Framingham, MA 01701, USA) at a flow rate of 40 μL/min for 2 minutes for each ionization mode. Lipid extracts were analyzed in both positive and negative ion modes for complete lipidome coverage using the TripleTOF 5600 System. Infusion MS/MSALL workflow experiments consisted of a TOF MS scan from m/z 200-1200 followed by a sequential acquisition of 1001 MS/MS spectra acquired from m/z 200 to 1200 (Simons et al., 2012). The total time required to obtain a comprehensive profile of the lipidome was approximately 10 minutes per sample. The data was acquired with high resolution (>30000) and high mass accuracy (~5 ppm RMS). Data processing using LipidView Software identified 150-300 lipid species, covering diverse lipids classes including major glycerophospholipids and sphingolipids. The peak intensities for each identified lipid, across all samples were normalized against an internal standard from same lipid class for the semi-quantitation purpose.

### Targeted Quantitation of Phosphatidylinositol (PI) and Lysophosphatidylinositol (LPI) Lipids

A targeted lipidomic assay for LPI and PI lipids was developed using HPLC on-line electrospray ionization tandem mass spectrometry (LC/ESI/MS/MS). Lipid extracts of cell lysates were prepared using a Folche extraction and normalized to total protein. *Standard Solutions:* The standards used in this assay were all purchased from Avanti Polar Lipids (LPI-16:0, LPI-18:0, LPI-18:1, LPI-20:4, PI-38:4). Internal standards used for these analyses were LPI-17:1, PI-34:1-d31; all of which were purchased from Avanti Polar Lipids. Standard LPI and PI species at concentrations of 0, 5, 20, 100, 500 and 2000 ng/ml were prepared in 90% methanol containing 2 internal standards at the concentration of 500 ng/ml. The volume of 5 μl was injected into the Shimadzu LCMS-8050 for generating the internal standard calibration curves. *HPLC Parameters:* A silica column (2.1 × 50mm, Luna Silica, 5 μm, Phenomenex) was used for the separation of PI and LPI species. Mobile phases were A (water containing 10 mM ammonium acetate) and B (acetonitrile containing 10 mM ammonium acetate). Mobile phase B at 95 % was used from 0 to 2 min at the flow rate of 0.3 ml/min and then a linear gradient from 95 % B to 50 % B from 2 to 8 min, kept at 50 % B from 8 to 16 min, 50 % B to 95 % B from 16 to 16.1 min, kept 95 % B from 16.1 to 24 min. *Mass Spectrometer Parameters:* The HPLC eluent was directly injected into a triple quadrupole mass spectrometer (Shimadzu LCMS-8050) and the analytes were ionized at ESI negative mode. Analytes were quantified using Selected Reaction Monitoring (SRM) and the SRM transitions (m/z) were 571 → 255 for LPI-16:0, 599 → 283 for LPI-18:0, 597 → 281 for LPI-18:1, 619 → 303 for LPI-20:4, 885 → 241 for PI-38:4, 583 → 267 for internal standard LPI-17:1, and 866 → 281 for internal standard PI-34:1-d31. *Data Analysis:* Software Labsolutions LCMS was used to get the peak area for both the internal standards and LPI and PI species. The internal standard calibration curves were used to calculate the concentration of LPI and PI species in the samples. All plasma LPI and PI species were normalized to the PI-34:1-d31 internal standard, while all tissue LPI species were normalized to the 17-1 LPI internal standard and all tissue PI species were normalized to the PI-34:1-d31 internal standard.

### RNA isolation and quantitative real time-PCR

Total RNA was isolated using an RNeasy or Trizol isolation methods using manufacturer’s recommendations (Qiagen and Thermo Fisher). RNA concentrations were quantified using a Nanodrop 2000. mRNA expression levels were calculated based on the ΔΔ-CT method. qPCR was conducted using the Applied Biosystems 7500 Real-Time PCR System. Primers used for qPCR are available on request.

### Cell Lines and Cell Culture Conditions

Caki-1 cell lines were obtained from American Type Culture Collection (ATCC), and were confirmed to be mycoplasma free. These cells were maintained in McCoy’s 5A Medium with 10% FBS and 1% Pen/Strep. Caki-1 expansions were propagated no longer than eighteen consecutive passages.

### Guide RNA Design and MBOAT7^−/−^ generation

The design of the MBOAT7 sgRNA was optimized using open source software CRISPOR (55). Synthetic sgRNA were designed to target all transcript variants of MBOAT7. Prior to nucleofection, the sgMBOAT7 at a working concentration of 30uM and recombinant Cas9 protein were incubated together at 6:1 ratio (sgMBOAT7:Cas9 protein) as recommended by manufacturer (Synthego). Parental Caki-1 cells were then nucleofected, and single cell isolated for clonal expansion. After growing and passaging cells from single cell suspension, multiple independent clonal populations were then screened for knockout populations. The chromatographs from the clonal populations were screened for editing using the previously described method (56). These populations were then validated using targeted mass spectrometry for LPI and PI lipids, mRNA abundance by qPCR, and Western blotting.

### Cell Viability Assay

At day zero, two thousand cells were seeded in 96-well plates. A relative cell number was determined using Cell Titer Glo® (Promega, Madison, WI, USA), according to manufacterer’s instructions. In this assay we used 1:1 buffer solution to growth media at endpoint. Then measure luminescence after 1 hour of incubation. Time points were performed for 48hrs, 96hrs, and 120hrs.

### Cell Cycle Analysis

Using a previously described method from BioLegend (Cat# 640914). In brief, we plated 35mm plates (n=4 per group) seeded at 200K cells for 2 days. These cells were then isolated, washed and fixed in 70% ethanol. These cells were then stained at room temperature in the dark in Propidium Iodide (Cat# 421301) and Annexin V for 15min. These cells were then analyzed using LSRFortessa (BD Biosciences). We used FlowJo software (Tree Star Inc.) to determine Propidium Iodide staining after excluding PI^+^Annexin V^+^ apoptotic cells.

### Migration Assay

The *in vitro* scratch was performed as previously described (57). Briefly, prior to seeding cells, 6-well plates were coated with an extracellular matrix. Approximately 200,000 cells were plated into each well in the 6-well plate. These wells grew to 90-100% confluency. After reaching confluency, plates were scratched and imaged for 16 hours every 10 minutes. Three separate experiments were conducted with 2 replicates per experiment. The quantification of these experiments was done using the scratch area of the first image and last image. Using ImageJ, the percent change in area was quantified using the following formula:

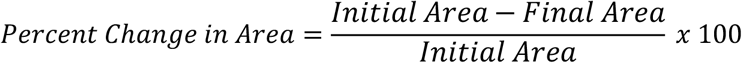

### In Vivo ccRCC Xenograft Studies

MBOAT7^+/+^ and MBOAT7^−/−^ Caki-1 cell lines were injected in the subcutaneous flank of NSG animals (Jackson laboratory) at 2.5 million cells per mouse in PBS (n=10 per group). Once tumors were palpable, digital caliper measurements were used to follow tumor growth over time. All mice studies were approved by the Institutional Animal Care and Use Committee of the Cleveland Clinic.

### Western Blot Analysis

Cell tissue lysates were generated using a modified RIPA buffer and western blotting was performed following the previously described methods (58).

### Statistical Analysis

All figures are shown as mean ± SE. For comparisons of three groups, we utilized one-way ANOVA with a *post hoc* Tukey test. Log-rank test (Mantel-Cox test) was used to compare survival differences between two groups of patients. When comparing two groups, we used an unpaired Student t test or multiple t test. The P values significance cutoffs for all tests used are as follows: p-value < 0.05(*), 0.002(**), 0.0002(***), <0.0001(****).

## Author Contributions

C.K.A.N., J.D.S., J.D.L., B.I.R., J.M.B. planned the project, designed experiments, and wrote the manuscript; C.K.A.N., R.Z., D.J.S., C.A.T., V.V. and D.B. executed experiments. C.K.A.N., R.Z., D.J.S., V.V. and D.B. analyzed the data; J.D.L., B.I.R., and J.M.B. provided financial support; all authors were involved in the editing of the final manuscript.

## Acknowledgments

We would like to thank the patients from Taussig Cancer Center of the Cleveland Clinic who donated their tumor samples for our study, and members of the Lathia laboratory for constructive directions on this research project. This study was supported in part by grants provided by the National Institutes of Health (NIH): R01 HL122283 (J.M.B.), P50 AA024333 (J.M.B.). Development of lipid mass spectrometry methods reported here were supported by generous pilot grants from the Clinical and Translational Science Collaborative of Cleveland (4UL1TR000439) from the National Center for Advancing Translational Sciences component of NIH and the NIH Roadmap for Medical Research, the Case Comprehensive Cancer Center (P30 CA043703), the VeloSano Foundation, and a Cleveland Clinic Research Center of Excellence Award. DB was supported by a Case Comprehensive Cancer Center training grant (T32CA059366)

