## Supplemental Figures for "MBOAT7-Driven Phosphatidylinositol Remodeling Promotes the Progression of Clear Cell Renal Carcinoma"

Supplemental File

Supplementary Fig. 1: Other Land Cycle Remodeling Pathways in ccRCC

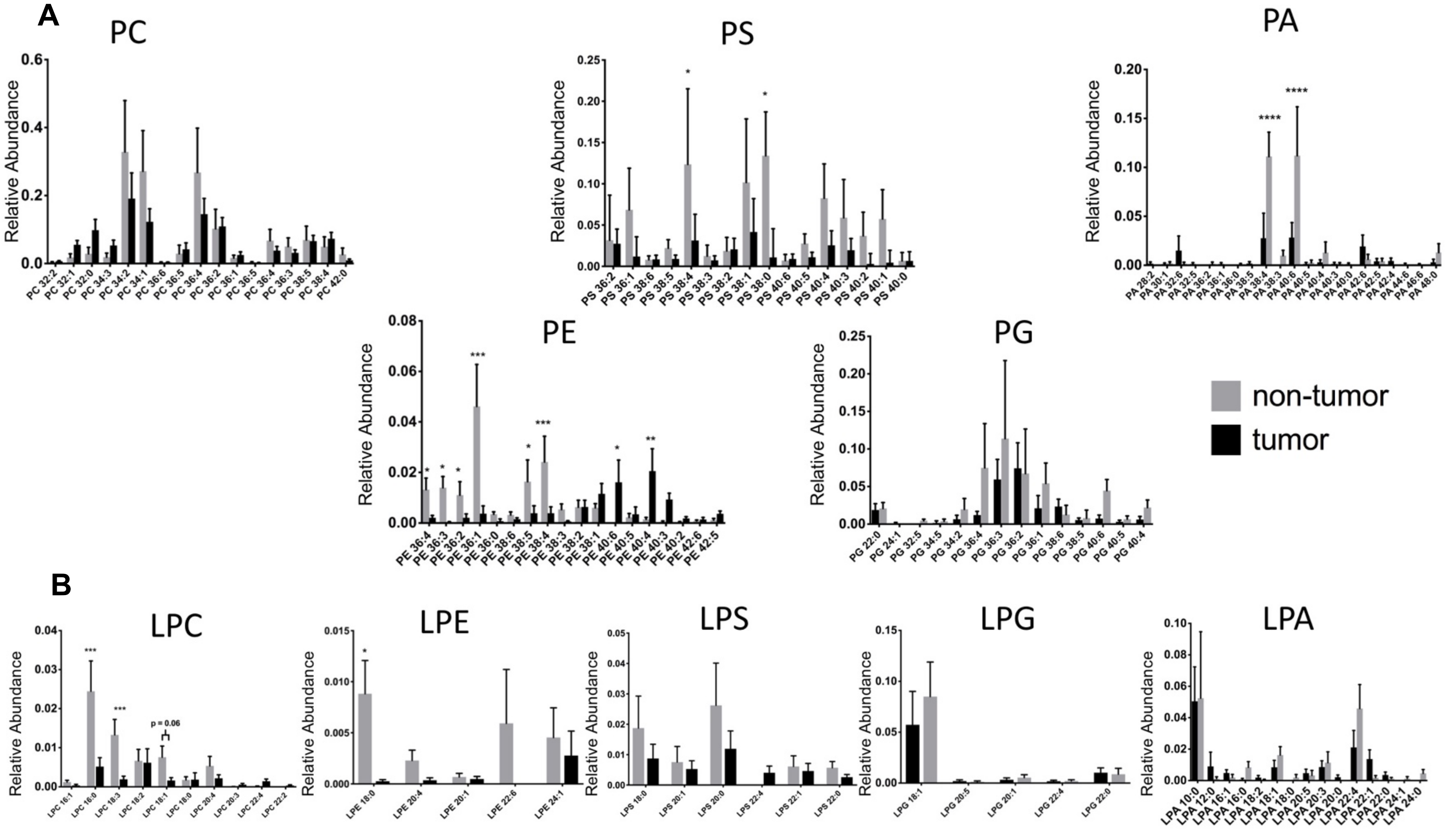


**Supplementary Figure 1: Other phospholipid changes in metastatic vs. non-tumor lipidomics**. A) Phosphatidylcholine, Phosphatidylethanolamine, phosphatidylserine, phosphatidylglycerol and phosphatidic acid in the tumor and non-tumor. B) Lyso-phosphatidylcholine, lysophosphatidylethanolamine, lysophosphatidylserine, lysophosphatidylglycerol, lysophosphatidic acid in the tumor and non-tumor. (n=10 per group) Student t-test and p-value: < 0.05(*), 0.0021(**), 0.0002(***), <0.0001(****).

Supplementary Fig. 2 – Sanger Sequencing confirms CRISPR knockout of MBOAT7

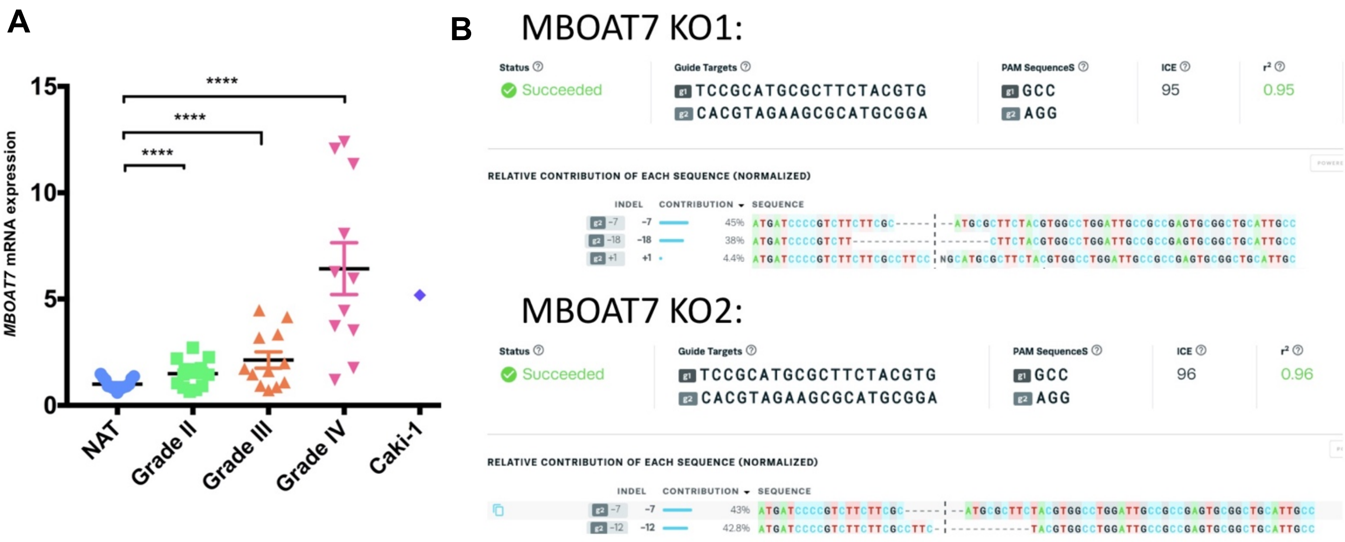


**Supplementary Figure 2: Sequencing validation of our MBOAT7 loss of function in Caki-1 cell model**. A) *MBOAT7* mRNA expression across tumor grade and the Caki-1 cell model (n= 1 per group). B) Sanger sequencing of the sgMBOAT7 cut site in two Caki-1 clones aligned to the Caki-1 wild-type sequence using ICE software. Student t-test and p-value: < 0.05(*), 0.0021(**), 0.0002(***), <0.0001(****).

Supplementary Fig. 3 - Free Arachidonic Acid Tracing Study into the phospholipid pools

**A**

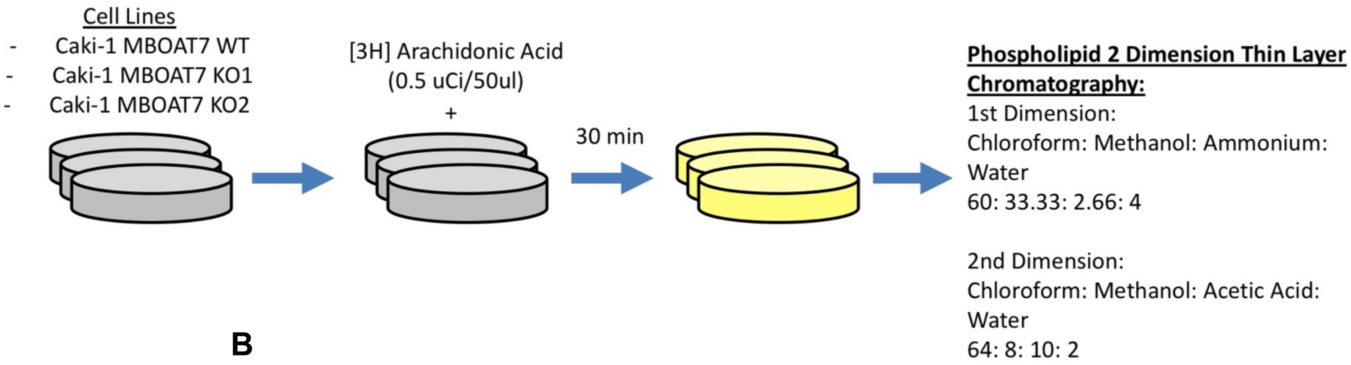


**B**

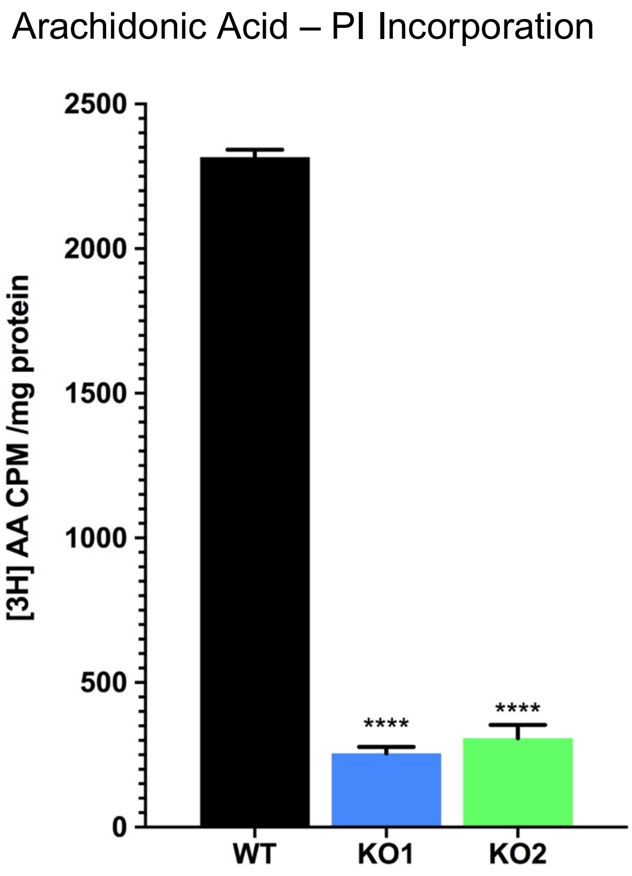


**Supplementary Figure 3: Radiolabeled tracing of $^{3}$H-Arachidonic Acid into Phospholipid pools**. A) Schematic of the radiolabeled arachidonic acid experimental design. B) 3H-Arachidonic acid (3H-AA) tracing into phospholipid pools using 2-Dimensional Thin-Layered Chromatography. Both knockouts compared to wild-type demonstrate a ~90% reduction in 3H-AA incorporation into PI. (n=3 per group)

Supplementary Fig. 4 – RNAseq plots between MBOAT7$^{+/+}$ and MBOAT7$^{-/-}$ clones

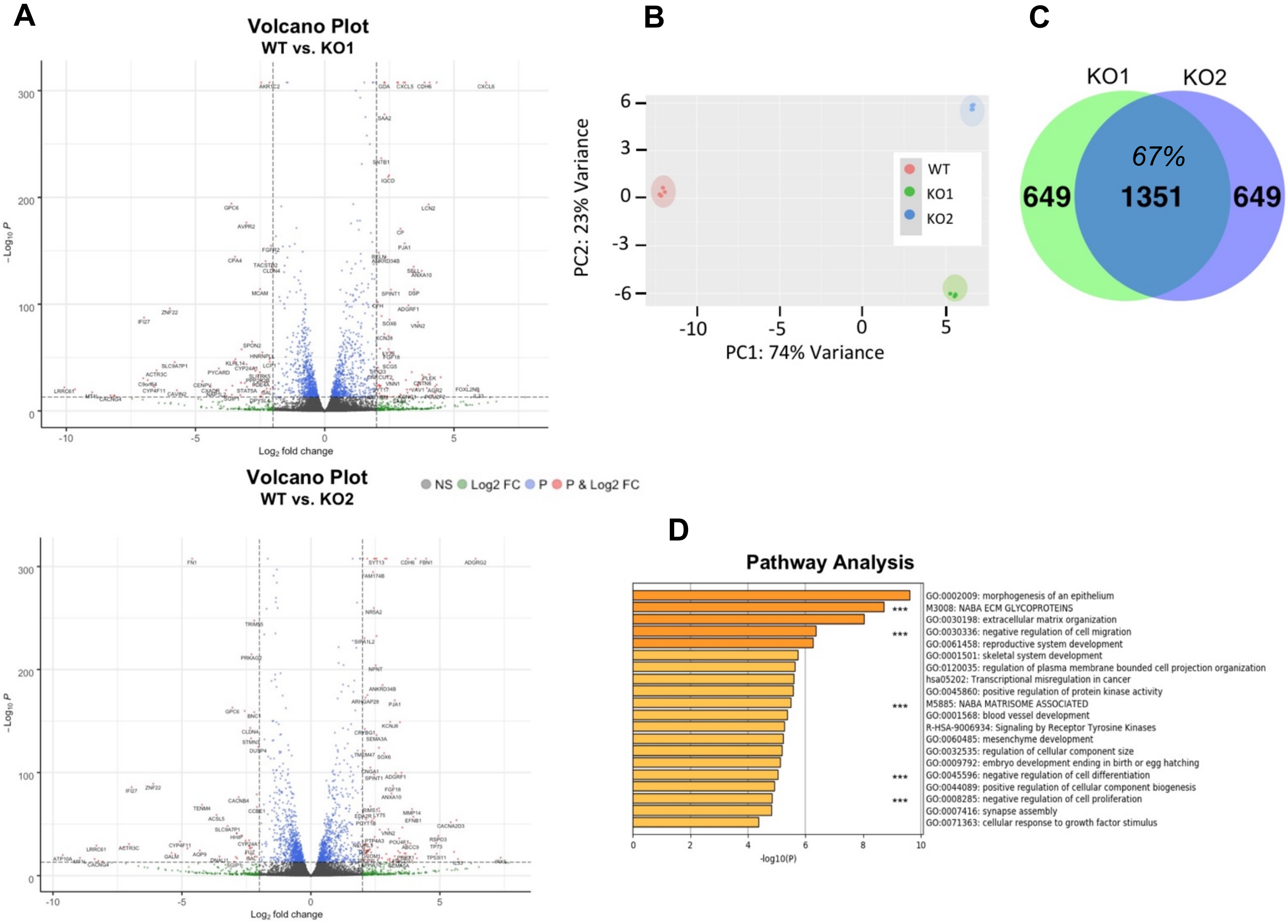


**Supplementary Figure 4: RNAseq plots comparing WT, KO1, and KO2. A)** Volcano Plot highlighting differentially regulated genes in KO1 and KO2 compared to WT (N=4 per group) **B)** PCA plot of the three groups: WT, KO1 and KO2. **D)** Pathway analysis for KO2 top 200 upregulated transcripts. ***< shared with KO1 pathway analysis in Figure 4.

**Supplementary Movie 1-3: *In vitro* migration movies comparing WT, KO1, and KO2 over a 16 hour period. A)** The in vitro scratch assay migration movies of WT, KO1, and KO2 over the same 16 hour time frame. (N= 2 per group per experiment) 3 independent experiments
